# Ephemeral-habitat colonization and neotropical species richness of *Caenorhabditis* nematodes

**DOI:** 10.1101/142190

**Authors:** Céline Ferrari, Romain Salle, Nicolas Callemeyn-Torre, Richard Jovelin, Asher D. Cutter, Christian Braendle

## Abstract

**Background:** The drivers of species co-existence in local communities are especially enigmatic for assemblages of morphologically cryptic species. Here we characterize the colonization dynamics and abundance of nine species of *Caenorhabditis* nematodes in neotropical French Guiana, the most speciose known assemblage of this genus, with resource use overlap and notoriously similar outward morphology despite deep genomic divergence.

**Methods:** To characterize the dynamics and specificity of colonization and exploitation of ephemeral resource patches, we conducted manipulative field experiments and the largest sampling effort to date for *Caenorhabditis* outside of Europe. This effort provides the first in-depth quantitative analysis of substrate specificity for *Caenorhabditis* in natural, unperturbed habitats.

**Results:** We amassed a total of 626 strain isolates from nine species of *Caenorhabditis* among 2865 substrate samples. With the two new species described here (*C. astrocarya* and *C. dolens*), we estimate that our sampling procedures will discover few additional species of these microbivorous animals in this tropical rainforest system. We demonstrate experimentally that the two most prevalent species (*C. nouraguensis* and *C. tropicalis*) rapidly colonize fresh resource patches, whereas at least one rarer species shows specialist micro-habitat fidelity.

**Discussion:** Despite the potential to colonize rapidly, these ephemeral patchy resources of rotting fruits and flowers are likely to often remain uncolonized by *Caenorhabditis* prior to their complete decay, implying dispersal-limited resource exploitation. We hypothesize that a combination of rapid colonization, high ephemerality of resource patches, and species heterogeneity in degree of specialization on micro-habitats and life histories enables dynamic co-existence of so many morphologically cryptic species of *Caenorhabditis*.

## Introduction

A long-standing question in ecology is how a diversity of species can coexist simultaneously in sympatry. A hotspot of *Caenorhabditis* nematode diversity in the tropical rainforest of French Guiana, in particular, provides a compelling example in which many species occupy the same micro-habitats [1]. This speciose genus (>50 species, K. Kiontke pers. comm.) exhibits frequent morphological crypsis and resource overlap among species, despite extreme genomic divergence between one another [2–5]. One possible contributor to regional species co-existence is resource patch turnover, especially when species differ in their colonization and competitive abilities [6–8].

*Caenorhabditis* nematodes thus provide a tantalizing ecological model for metapopulation dynamics as a contributor to species co-existence, in addition to their role as important biomedical model organisms [9]. All species of *Caenorhabditis* are microbivorous, found most commonly within decaying vegetal matter. These ephemeral resource patches should underlie a highly dynamic metapopulation structure for *Caenorhabditis* on the landscape, especially in tropical environments with rapid decay rates [10–13]. However, among the many poorly understood features of their natural history, the details of patch colonization, occurrence with patch age, and patch substrate specificity form especially crucial factors for understanding the abundance and diversity of these organisms.

Natural *Caenorhabditis* populations have become increasingly studied during the past few years, with a primary focus on *C. elegans* [2, 12–18]. These efforts have led to the discovery of many new species and the establishment of hundreds of cryopreserved *Caenorhabditis* wild isolates available for study from across the world [3, 10, 11, 19]. In turn, these live specimen collections have prompted studies of population genomics [20, 21], genome-wide association mapping of biologically important traits [22], discovery of and experimentation with natural pathogens [11, 23], and the seasonal turnover of species [16]. Together they provide the unique opportunity to integrate the extensive knowledge of *Caenorhabditis* cellular development, neurobiology, and molecular genetics with ecological characteristics and evolutionary processes.

In-depth analyses of natural *Caenorhabditis* populations, however, have been limited to mainland France and Germany, with a focus on the three predominant species of this region: *C. elegans, C. briggsae* and *C. remanei* [10–12, 15–17, 24–26]. The vast majority of these surveys were collected in anthropogenic habitats, such as orchards or garden compost heaps, making analysis of habitats unperturbed by extensive human activity crucial to understanding the diversity and abundance of these organisms. Documentation of natural populations in tropical regions is less secure, despite providing epicenters of biodiversity from the levels of genetic variability to species richness [1, 10, 27]. Previous study of *Caenorhabditis* from tropical French Guiana brought more general ecological questions to the fore [1, 27]. After having identified the most locally-common species, this work demonstrated co-occurrence of distinct species and genotypes within individual micro-habitat patches, and quantified DNA sequence variability for the two self-fertilizing species found there [1, 27]. Consequently, quantitative analysis of species richness and abundance, species-specificity of micro-habitat substrates, phoretic associations, and dispersal dynamics represent essential outstanding problems.

To characterize the dynamics and specificity of colonization and exploitation of ephemeral resource patches, we conducted manipulative field experiments and the largest sampling effort to date for *Caenorhabditis* outside of Europe [12, 15–17, 24–26]. By quantifying colonization rates and distinct species incidence among micro-habitats, we provide a foundational view of key factors in the metapopulation dynamics that we predict to be important in enabling local and regional species co-existence.

## Methods

### Collection of samples and analysis

We collected samples during field expeditions in April 2013, August 2014 and August 2015. Most samples were collected in primary rainforest of the Nouragues Natural Reserve (4°5’ N, 52°4’ W), close to the two CNRS field stations “Inselberg” and “Parare,” 8km apart. In addition, we conducted opportunistic sampling of rotting fruit/flower substrates from other sites in French Guiana, mostly in primary and secondary forests of coastal regions (2013, 2015) and in the surroundings of the village Saül in central French Guiana (2015) (Figure 1). Additional File 1 lists all strain isolates collected, with corresponding details on sampling sites and substrate characteristics. All isolates were cryopreserved and are available upon request.

**Figure 1.**
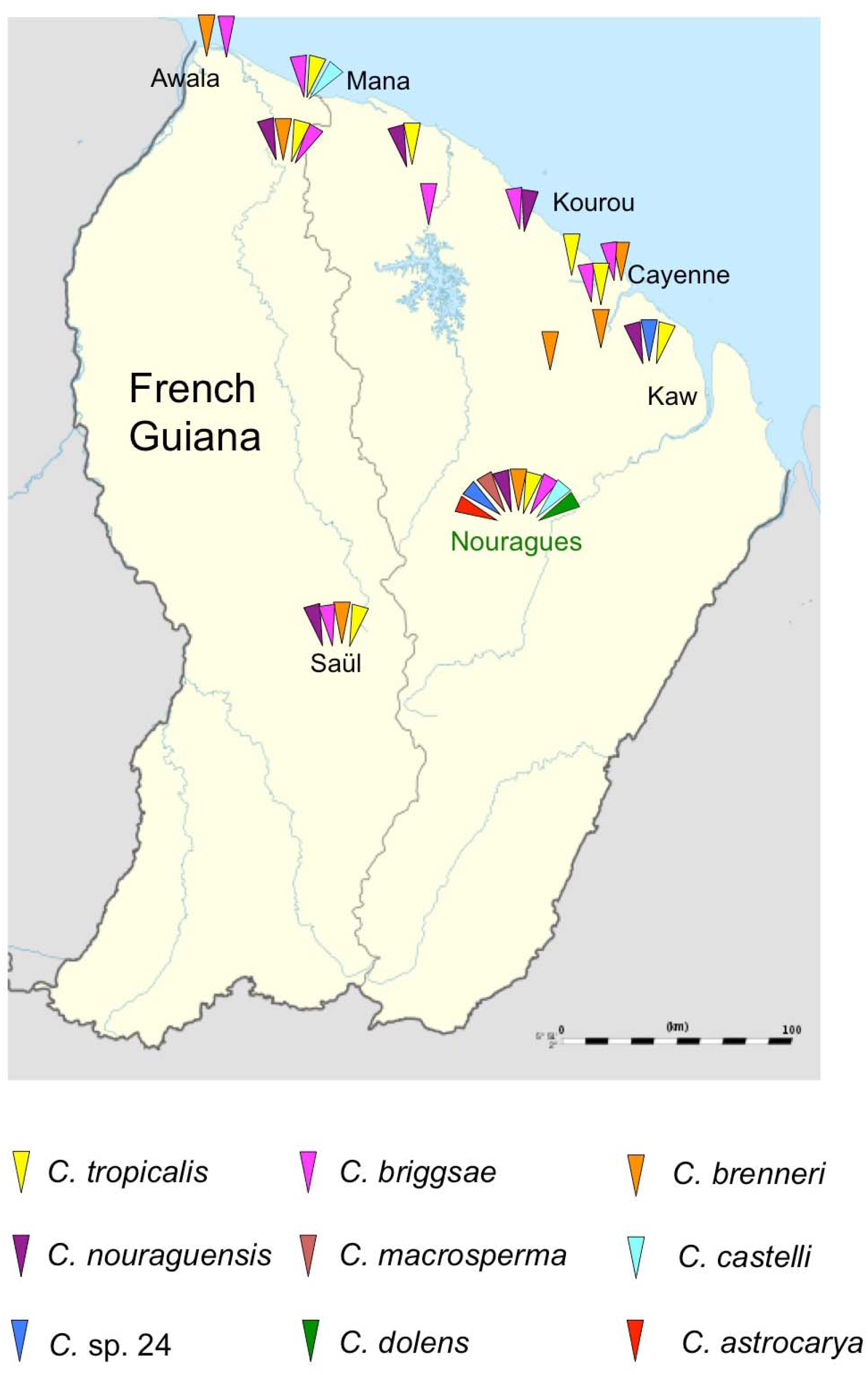
Overview of *Caenorhabditis* species distribution in French Guiana (2009 – 2015), including data of Félix et al. (2013).

### Identification of species, isolate establishment, and species richness estimation

For substrate sampling, nematode isolation, and species identification, we followed previously established protocols [1, 28, 29]. In brief, we stored samples in sealed plastic bags prior to analysis in the laboratory; a small subset of samples was directly isolated on site. Based on measurements of substrate samples from collections in 2013, they weighed ~20g on average and ranged from ~5g to ~100g. To isolate nematodes, substrate samples were placed on Nematode Growth Medium (NGM) plates inoculated with the *E. coli* strain OP50 [30]. This procedure allows efficient isolation of a *Caenorhabditis* species from each sample, but is biased against identifying multiple species, in particular, slow-growing species. *Caenorhabditis* genus identity was first established through microscopy analysis of morphology [31], and species identity was then confirmed using sequencing of the ITS2 region (Internally Transcribed Spacer) between the 5.8S and 28S rDNA genes, as previously described [2, 3]. We used primers 5.8S-1 (5’-CTGCGTTACTTACCACGAATTGCARAC) and KK28S-4 (5’-GCGGTATTTGCTACTACCAYYAMGATCTGC) that amplify a fragment of approximately 2kb, which was then partially sequenced using the forward PCR primer and the sequencing primer KK-28S-22 (5-CACTTTCAAGCAACCCGAC) [3, 31]. The sequence of the ITS2 fragment provides a reliable species identification tag as it is highly divergent between, but not within, *Caenorhabditis* species [3]. Partial ITS2 sequences of the two new species (*C. astrocarya* and *C. dolens*) will be deposited in GenBank (Additional File 9).

All isolates were derived from a single hermaphrodite at the larval stage (for androdioecious species) or an adult, mated female (for gonochoristic species). All isolates were then cryopreserved (see Additional File 1 for complete list of isolates).

We estimated total species richness using the Chao2 estimator with EstimateS v. 9.1.0 [32].

### Identification and description of novel species

We provide descriptions for two new species, which were provisionally termed *C*. sp. 37 and *C*. sp. 42. We followed the current standard for new species description in this group based on molecular barcodes and biological species inference from mating tests [2]. ITS2 barcode sequences for both species are highly distinct from all currently known *Caenorhabditis* species, supporting their new species status. Nevertheless, following previously described protocols [3], we carried out reciprocal mating crosses with species that show strongest ITS2 sequence similarity to affirm reproductive isolation and identity as distinct biological species (see below).

### Nomenclatural acts

The electronic edition of this article conforms to the requirements of the amended International Code of Zoological Nomenclature, and hence the new names contained herein are available under that Code from the electronic edition of this article. This published work and the nomenclatural acts it contains have been registered in ZooBank, the online registration system for the ICZN. The ZooBank LSIDs (Life Science Identifiers) can be resolved and the associated information viewed through any standard web browser by appending the LSID to the prefix “http://zoobank.org/”. The LSID for this publication is urn:lsid:zoobank.org:pub:06DBDCA0-DD60-43BA-9A99-14F0748D4FE2. The electronic edition of this work was published in a journal with an ISSN.

### Substrate incidence of *Caenorhabditis*

To obtain quantitative estimates of *Caenorhabditis* presence on different types of substrates (native fruit versus litter), we sampled 56 sites across the trail network of the Inselberg station in the Nouragues Natural Reserve (2013) (Figure 2, Additional File 2). At each site, we defined a sampling area of approximately 20m^2^ within which we collected ~10 litter samples and up to five fruit samples, the latter consisting of diverse, usually strongly decayed fruit material on the forest floor.

**Figure 2.**
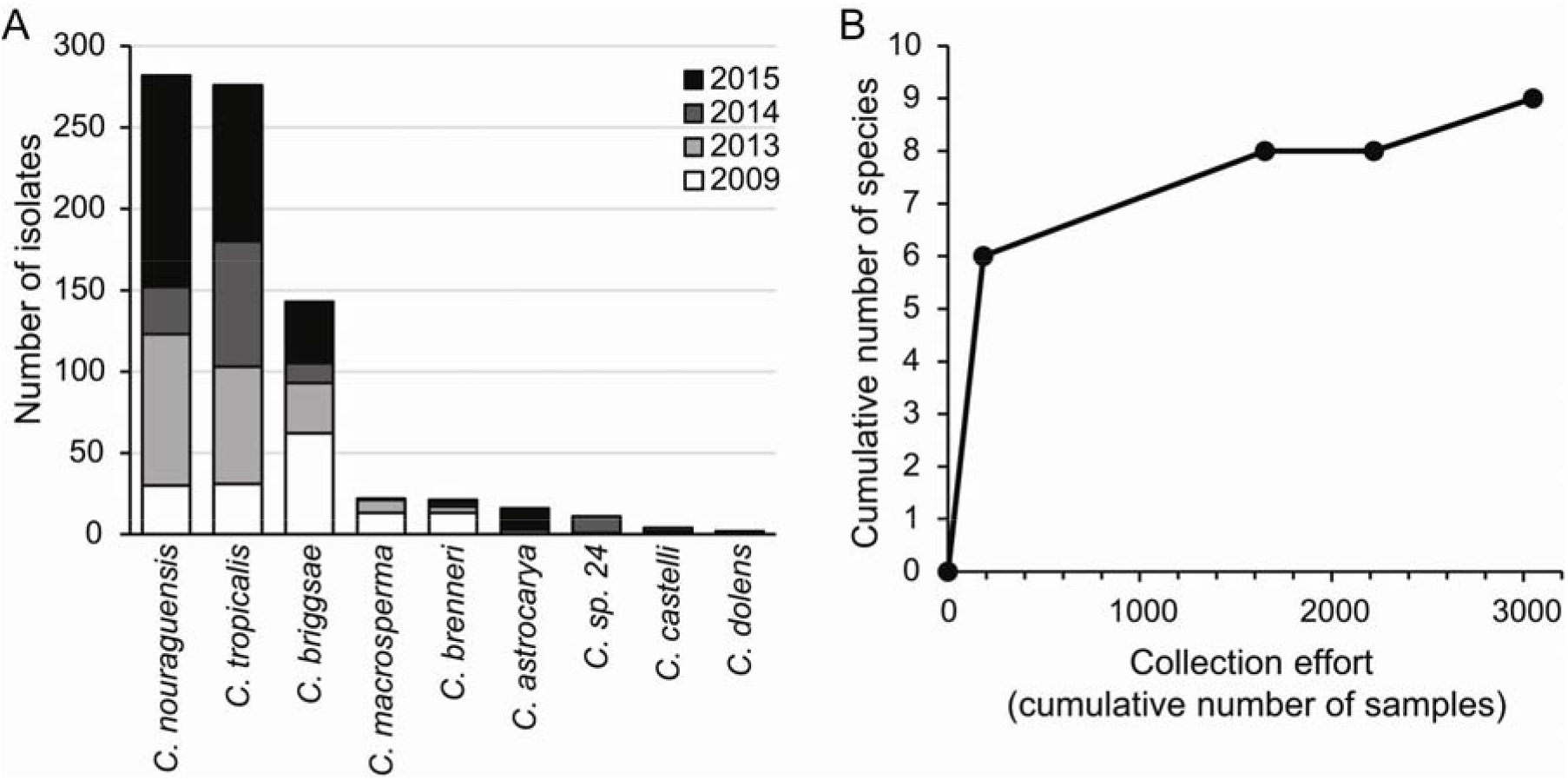
Species diversity and abundance of *Caenorhabditis* nematodes collected in French Guiana. **(A)** Species rank-abundance distribution for *Caenorhabditis* isolates collected from 2009 to 2015. **(B)** Collectors curve of species identified in French Guiana as a function of sampling effort from 2009 to 2015.

### Micro-habitat colonization and age-dependent incidence

To assess *Caenorhabditis* colonization of bait samples (Additional File 3), we distributed pieces of surface-sterilized oranges (each piece 1/16^th^ of a large orange) at 72 intervals spaced approximately 30m apart along the trail network of the Parare station in the Nouragues Natural Reserve (2014). At each spot, we distributed three groups of three orange pieces on the ground within an area of approximately 3m^2^, with a single sample representing a group of three orange pieces, which were placed together into a single plastic bag four days after. Therefore, we collected three independent bait samples at each spot (total samples: N=216).

To assess *Caenorhabditis* colonization rates of bait samples at localities that had abundant rotting fruit/flower substrates on the ground (Additional File 4), we selected five localities near the Parare station and 18 localities near the Inselberg station (Nouragues Natural Reserve). At each locality, we distributed five bait samples (orange and banana pieces) within an area of approximately 5m^2^. After four days, bait samples as well as five native fruit/flower samples were collected and stored in plastic bags for further analysis.

To examine *Caenorhabditis* colonization as a function of substrate age, we focused on a single site where *Clusia* flowers (likely *Clusia palmicida*) at different stages of decay covered the ground (Inselberg station, Nouragues Natural Reserve). We haphazardly collected 70 flower samples across the entire ~75m^2^ site from each of three clearly distinct, progressive stages of decay (N=210; Figure 4, Additional File 5). The freshest decay stage comprised fleshy petals and intact floral resin [33], with the latest stage being composed primarily of fibrous plant tissue. We then assessed species incidence on each flower sample. In addition, for a subset of these samples containing *Caenorhabditis* (N=21), we immediately placed samples on NGM plates seeded with *E. coli* and determined number and stages of individuals five to seven hours later (Additional File 6).

## Results

Through the combined effort of targeted sampling of native substrates, experimental baits, and opportunistic collection in the Nouragues Natural Reserve and throughout French Guiana across three years, we amassed a total of 626 strain isolates from nine species of *Caenorhabditis* among 2865 substrate samples (Figures 1 and 2A, Additional File 1). This represents the largest collection effort for *Caenorhabditis* to date outside of mainland Europe, and the single largest collection effort for non-anthropogenically modified habitats. Integrating the sampling and species discovery efforts in this study with previous collection leads to a species richness upper 95% confidence interval bound of 9.87 *Caenorhabditis* in French Guiana (Chao2 estimator), suggesting that at most one additional species is likely to be discovered given current sampling procedures (Figure 2B). Three species provide first records in French Guiana and two of these species are new to science (*Caenorhabditis dolens* n. sp. = *C*. sp. 37 and *Caenorhabditis astrocarya* n. sp. = *C*. sp. 42, see below). Four species were most widespread by being present at numerous inland and coastal localities (*C. nouraguensis*, *C. tropicalis*, *C. briggsae*, and *C. brenneri*), with the gonochoristic *C. nouraguensis* and the androdioecious *C. tropicalis* being most abundant (Figure 2A). Thus, French Guiana harbours a total of five endemic species to date (*C. nouraguensis, C. macrosperma, C. castelli, C. dolens* n. sp. = *C*. sp. 37 and *C. astrocarya* n. sp. = *C*. sp. 42) in addition to the presence of three cosmopolitan species (*C. tropicalis, C. briggsae, C. brenneri*) and one species found previously elsewhere in the New World tropics (C. sp. 24 found in Panama, M. Rockman, pers. comm.).

### Substrate incidence of Caenorhabditis

We first aimed to test for quantitative differences in the prevalence of *Caenorhabditis* between decaying vegetal micro-habitats within a given site, namely between leaf litter and rotting fruit. Across 560 leaf litter samples from 56 locations covering about 4 km^2^ (Figure 3), we isolated only two species of *Caenorhabditis* from just 2.7% of the samples, with 43 of the 56 locations lacking *Caenorhabditis* entirely in litter samples (Additional File 2). By contrast, we isolated three *Caenorhabditis* species among 20.6% of 209 rotting fruit samples from these same locations, failing to find representatives of the genus from just 20 of the locations (Fisher’s Exact Test P<0.0001) (Additional File 2). Just three of the locations contained *Caenorhabditis* in litter samples only, in contrast to 17 of the 56 locations that had *Caenorhabditis* found exclusively in fruit samples (Figure 3). These results provide quantitative support for the received wisdom that *Caenorhabditis* occurs more frequently in high-nutrient rotting micro-habitats [10, 11, 14]. We isolated multiple species of *Caenorhabditis* at six of the 56 sampling locations (Figure 3). Non-*Caenorhabditis* nematodes (predominantly species of the Rhabditidae and Diplogasteridae) were found in most litter samples (98%) and fruit samples (88%) Additional File 2).

**Figure 3.**
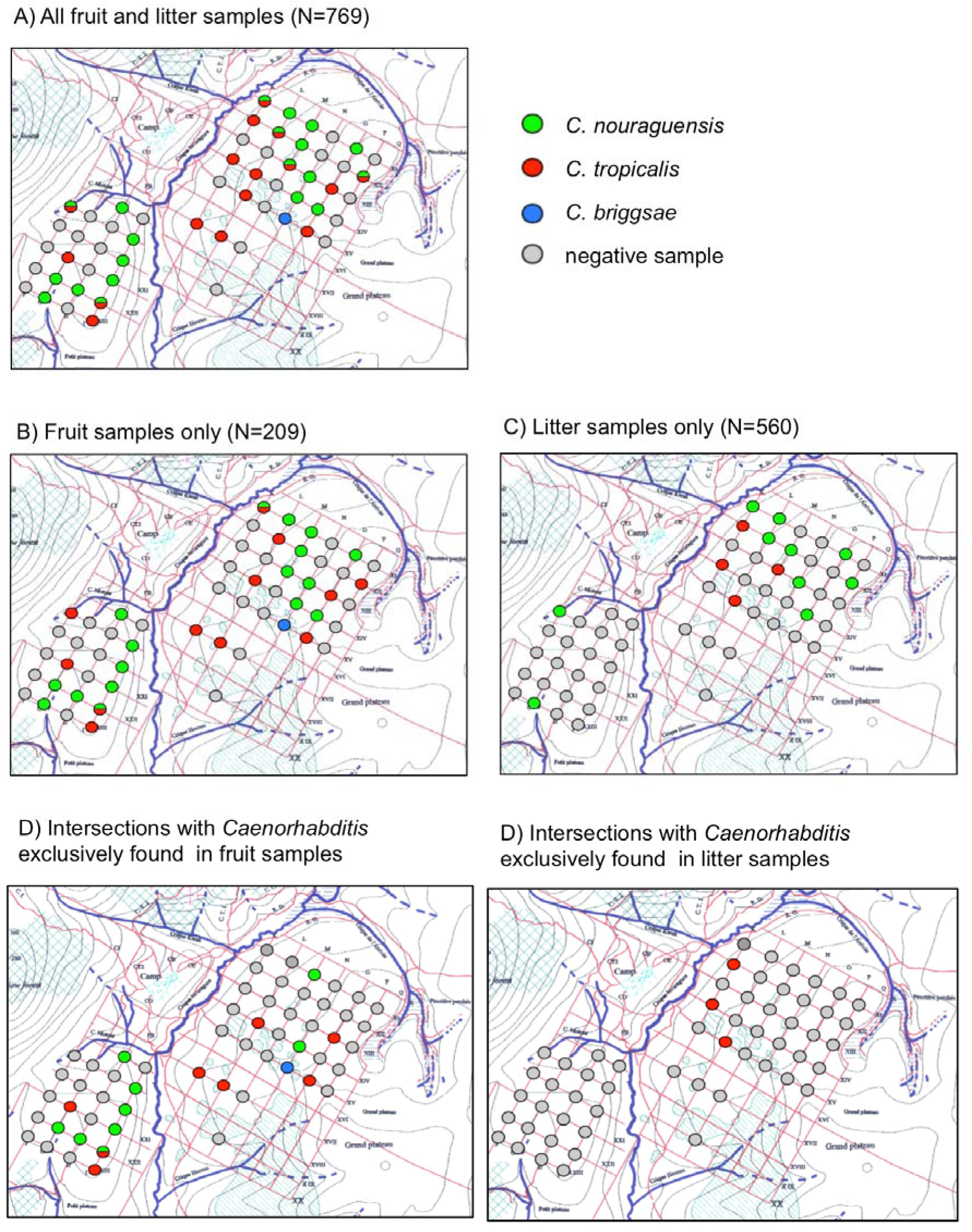
Substrate incidence of *Caenorhabditis* (Inselberg site, Nouragues Natural Reserve). Quantitative differences in the incidence of *Caenorhabditis* in decaying leaf litter versus rotting fruit. Red grid lines in sampling area spaced ~100m apart.

### Micro-habitat colonization and substrate age-dependent incidence

We next performed a series of field experiments to quantify colonization of fresh substrates by *Caenorhabditis*. First, we placed three sterilized orange fruit baits at each of 72 sites along multiple transects near the Parare Station within the Nouragues Natural Reserve, and collected them after four days to test for colonization. Overall, we isolated *Caenorhabditis* on 9.7% of a total of 216 bait samples (*C. tropicalis, C. briggsae, C. nouraguensis*), with a range of 4% to 44% of baits having been colonized across the transects (Figure S1 and Additional File 3). We isolated two species (*C. nouraguensis* and *C. tropicalis*) from one bait sample, although this is an underestimate of the coincidence of species because our isolation protocol is biased against finding multiple species per sample. Of note, only a single bait sample was colonized by a non-*Caenorhabditis* nematode. These findings establish that *Caenorhabditis* colonizes new nutrient-rich substrates rapidly and more efficiently than do other kinds of nematodes. Nevertheless, given the observed colonization rates of fresh baits, many resource patches are likely to remain uncolonized by *Caenorhabditis* before complete fruit decay. As a result, resource exploitation by *Caenorhabditis* is likely to be limited by dispersal to fresh patches.

Having established that *Caenorhabditis* occurs readily on both native substrates of unknown age and on fresh baits, we next tested whether fresh fruit baits got more efficiently colonized by *Caenorhabditis* (*C. tropicalis, C. briggsae, C. nouraguensis, C*. sp. 24) when they were positioned near to locations on the rainforest floor with a high natural density of rotting fruit or flowers compared to random locations. Indeed, 29.8% of baits (N=124 from a total of 23 sites) near to high densities of rotting native fruit or flowers contained *Caenorhabditis* after 5 days (Additional File 4), compared to just 9.7% of randomly-located bait samples (N=216) (Fisher’s Exact Test, P<0.0001) (see above; Additional File 3). Baits and native fruits in the same vicinity had a similar incidence of *Caenorhabditis* (29.8% on baits versus 20.0% on native fruits) (Fisher’s Exact Test, P=0.10). In contrast, Non-*Caenorhabditis* nematodes were more common on the native fruit/flower substrates (58 of 115 baits colonized) than on the fresh baits (27 of 124 baits colonized) (Fisher’s exact test, P<0.0001; Additional File 4). These findings implicate existing rotting fruit/flower as an important dispersal source for *Caenorhabditis* in colonizing new micro-habitats.

As a final means of characterizing the incidence of *Caenorhabditis* as a function of substrate age, we quantified nematode species among 210 *Clusia* flowers in three distinct phases of decay (Figure 4, Additional File 5). We found a striking effect of substrate decay stage on the presence of *C. nouraguensis:* more than half of the 70 freshest samples had been colonized (54%), but just 13% and 4% of samples in increasingly more advanced states of decay contained this species (Figure 4). *C. tropicalis* was rarer in these samples overall, but was never found on those *Clusia* samples in the most-advanced stage of decay (Figure 4). By contrast, non-*Caenorhabditis* nematodes were least common on the freshest samples of *Clusia* (Figure 4). Characterizing a subset of *Clusia* flowers containing *Caenorhabditis* (N=21) immediately on site, we found that most of these positive samples were colonized by starved adults rather than juvenile or dauer stages (Additional File 6). Only 3 of 21 samples contained more than 40 individuals of mixed age classes, indicative of proliferation, suggesting that colonization of fresh resource patches get founded by very few individuals. This low incidence of proliferating nematode populations contrasts with *C. elegans* on rotting apples in orchards [18].

**Figure 4.**
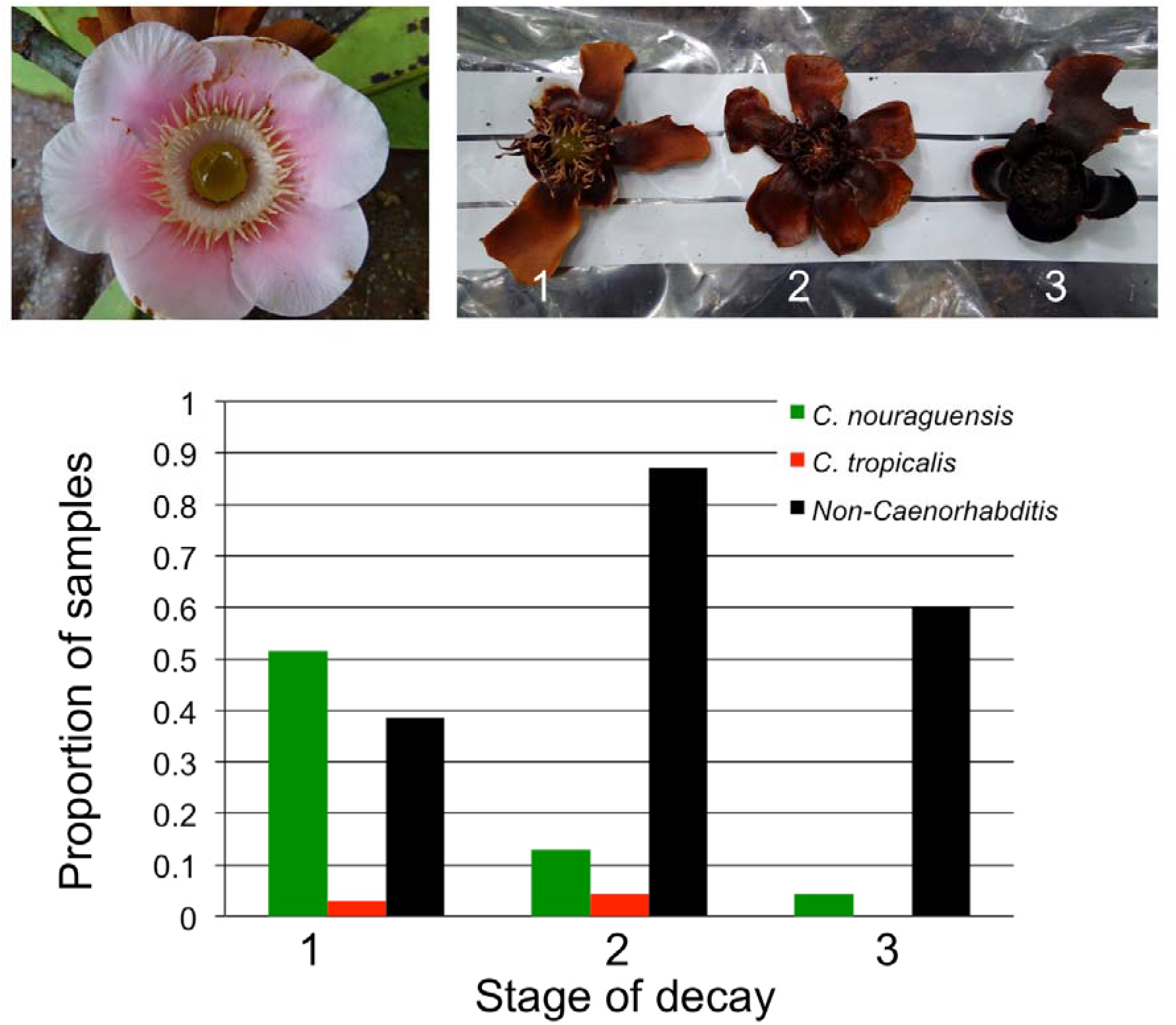
*Caenorhabditis* colonization of *Clusia* flowers at three distinct stages of decay. (1: slight decay, brown colour with intact floral resin, 2: intermdiate decay, brown colour with no floral resin; 3: strong decay, black colour with no floral resin). Incidence of nematodes based on sampling 70 flowers from each decay stage. Decay stage differs significantly for presence of *C. nouraguensis* (Fisher’s Exact Test P<0.0001) and non-*Caenorhabiditis* (Fisher’s Exact Test P<0.0001). Image on top-left shows a fresh *Clusia* flower.

### Landscape-scale assessment of neotropical species richness for *Caenorhabditis*

We complemented targeted and experimental characterization of *Caenorhabditis* substrates with opportunistic sampling of diverse ephemeral-habitat substrates throughout the Nouragues Natural Reserve (Figures S2 and S3, Additional File 7) and French Guiana more broadly (Figure 1 and Additional File 8). The more diverse substrate collections in the Nouragues Natural Reserve yielded isolation of five rarer species from 690 samples, in addition to the four more common species found in our targeted substrate sampling (as detailed in the previous section) (Additional File 7). Outside of the Nouragues Natural Reserve, we recovered 151 isolates from six species of *Caenorhabditis*, deriving from 210 samples from a locality in inland Saül and 247 samples from multiple localities along the Atlantic coast of French Guiana (Additional File 8). These six species comprise a subset of those nine *Caenorhabditis* found within the Nouragues Natural Reserve (Figure 1, Table 1), indicating that the species encountered throughout the regional scale can also be found from intensive sampling of a single locality.

**Table 1.** Overview of *Caenorhabditis* species and isolates collected from different localities in French Guiana (2013 – 2015). (A) Opportunistic sampling (2013, 2015) in French Guiana (excluding Nouragues). (B) Samples collected in the Nouragues Natural Reserve (2013, 2014, 2015).

| Locality | Year | Positive samples/Total | *C. astrocarya* | *C. brenneri* | *C. briggsae* | *C. castelli* | *C. dolens* | *C. macrosperma* | *C. nouraguensis* | *C. tropicalis* | *C. sp. 24* |
| --- | --- | --- | --- | --- | --- | --- | --- | --- | --- | --- | --- |
| Kourou | 2013 | 6/25 | 0 | 0 | 5 | 0 | 0 | 0 | 1 | 0 | 0 |
| Sentier St. Elie | 2013 | 7/27 | 0 | 0 | 0 | 0 | 0 | 0 | 4 | 3 | 0 |
| Angoulême | 2013 | 21/93 | 0 | 1 | 4 | 0 | 0 | 0 | 4 | 12 | 0 |
| Mana | 2013 | 6/25 | 0 | 0 | 2 | 1 | 0 | 0 | 0 | 3 | 0 |
| Awala-Yalimapo | 2013 | 10/19 | 0 | 2 | 8 | 0 | 0 | 0 | 0 | 0 | 0 |
| Macouria | 2013 | 1/4 | 0 | 0 | 0 | 0 | 0 | 0 | 0 | 1 | 0 |
| Saul | 2015 | 71/210 | 0 | 2 | 22 | 0 | 0 | 0 | 10 | 37 | 0 |
| Kaw mountains | 2015 | 25/45 | 0 | 0 | 0 | 0 | 1 | 0 | 14 | 10 | 0 |
| Matoury | 2015 | 4/9 | 0 | 0 | 3 | 0 | 0 | 0 | 0 | 1 | 0 |

B
| Locality | Year | Samples (total) | *C. astrocarya* | *C. brenneri* | *C. briggsae* | *C. castelli* | *C. dolens* | *C. macrosperma* | *C. nouraguensis* | *C. tropicalis* | *C. sp. 24* |
| --- | --- | --- | --- | --- | --- | --- | --- | --- | --- | --- | --- |
| Nouragues | 2013 | 161/1276 | 0 | 1 | 12 | 1 | 1 | 8 | 84 | 53 | 1 |
| Nouragues | 2014 | 134/567 | 3 | 2 | 12 | 0 | 0 | 1 | 29 | 77 | 10 |
| Nouragues | 2015 | 177/565 | 13 | 0 | 13 | 0 | 0 | 0 | 106 | 48 | 0 |

All three of the most common species were isolated in every collection period (2013, 2014, 2015), including a previous sampling effort in 2009 [1] (Figure 2A). Each of the remaining rarer species failed to be sampled in at least one collection episode, with the exception of *C. brenneri*, which was sampled in all collection periods in more human-associated locations. Our opportunistic sampling yielded 7 *Caenorhabditis* isolates on 58 invertebrates (12% incidence, found on insects and millipedes; Additional File 1); more detailed investigation of the potential for phoretic associations with tropical invertebrates is required in the future.

Estimation of total *Caenorhabditis* species richness from our sampling collector’s curve suggests that it is unlikely that more than one additional species will be recovered using current sampling procedures. The “Chao2” richness estimator yields an upper 95% confidence bound of 9.87 total species, compared to the 9 discovered in French Guiana to date. The species rank-abundance distribution for *Caenorhabditis* in French Guiana follows the typical pattern for low evenness of a few common species and many more rare species (Figure 2A), though no other species rank-abundance curves are available for comparison with *Caenorhabditis* in other regions for comparison.

### Discovery and description of two new species of *Caenorhabditis*

Our collection efforts yielded isolation of two species of *Caenorhabditis* previously unknown to science. Here we provide species descriptions of them, following the current standard for new species description in this group based on molecular barcodes and biological species inference from mating crosses [2].

*Caenorhabditis dolens* Braendle & Cutter sp. nov. urn:lsid:zoobank.org:act:4D39DD5D-2120-42E5-8521-0F35D8710592 = *Caenorhabditis* sp. 37 The type isolate by present designation is NIC394. The species is delineated and diagnosed by the fertile cross with the type isolate NIC394 in both cross directions, yielding fertile hybrid females and males that are interfertile and cross-fertile with their parent strains. The species reproduces through males and females. This species differs by ITS2 DNA sequences from all other species in [3] and [2] (Additional File 9). Note that these DNA sequences may vary within the species. From molecular data, it falls into the Drosophilae supergroup of *Caenorhabditis* with the closest species being *C*. sp. 38 (strain NIC559 from Dominica) and *C*. sp. 47 (strain NIC1127 from Peru), with which it does not form any larval progeny. The type isolate was collected in the Nouragues Natural Reserve (French Guiana) in 2013 from decaying fruit; no other isolates are currently known. The type culture specimens are deposited at the Caenorhabditis Genetics Center (<u>http://www.cbs.umn.edu/research/resources/cgc</u>).

*Caenorhabditis astrocarya* Braendle & Cutter sp. nov. urn:lsid:zoobank.org:act:EBE4FF82-F906-4237-9290-C84431F17D7D = *Caenorhabditis* sp. 42 The type isolate by present designation is NIC1040. The species is delineated and diagnosed by the fertile cross with the type isolate NIC1040 in both cross directions, yielding highly fertile hybrid females and males that are interfertile and cross-fertile with their parent strains. The species reproduces through males and females. This species differs by ITS2 DNA sequences from all other species in [3] and [2] (Additional File 9). Note that these DNA sequences may vary within the species. From molecular data, it falls into the Drosophilae supergroup of *Caenorhabditis* with the closest species being *C*. sp. 45 (strain NIC759 from Dominica) and *C. guadeloupensis* (strain NIC113 from Guadeloupe), with which it does not form any larval progeny. The type isolate was collected from pollen stalks of *Astrocaryum* palms (likely *Astrocaryum paramaca*) in the Nouragues Natural Reserve (French Guiana) in 2014 (Figure S4). Additional isolates (Additional File 1) were collected in the same location, exclusively from fresh and rotten pollen stalks on *Astrocaryum* plants in 2014 and 2015. The pollen substrate was coinhabited by abundant weevils. The type culture specimens are deposited at the Caenorhabditis Genetics Center (<u>http://www.cbs.umn.edu/research/resources/cgc</u>).

## Discussion

Deciphering the drivers of biodiversity and species coexistence first requires characterization of species richness and abundance, as well as detailed understanding of resource use and dispersal. To these ends for *Caenorhabditis* nematodes, a group most famous otherwise as biomedical model organisms, we conducted the largest biodiversity survey and field quantification of resource patch colonization in habitat unperturbed by human influence. Combining opportunistic sampling with quantitative, experimental methods, we collected nearly 3000 substrate samples and identified nine species of *Caenorhabditis* in French Guiana, two species of which are new to science (*C. astrocarya* n. sp. and *C. dolens* n. sp.). These species are roughly evenly divided among the three major phylogenetic subgroups within the genus (2 *Japonica* group species, 3 *Elegans* group species, 4 *Drosophilae* supergroup species) [3, 10]. Given current sampling procedures, we estimate that we are unlikely to discover more than one new additional species in this locality, which represents the greatest biodiversity hotspot for *Caenorhabditis* described to date.

The three most common species overall also were the most common species in every sampling year, and include both endemic (*C. nouraguensis*), regionally widespread (*C. tropicalis*), and globally distributed species (*C. briggsae*). Four of the rarer species are endemic to French Guiana given current neotropical sampling, but *C*. sp. 24 and *C. brenneri* are found elsewhere in tropical latitudes [34, 35] (M. Rockman, pers. comm.). The species identified at the regional scale were a subset of those found within the more-intensely sampled Nouragues Natural Reserve locality. In only one instance can we point to clear substrate specificity: *C. astrocarya* n. sp. was isolated exclusively in the rotting pollen of *Astrocaryum* palms, on which it frequently occurs. We also note that *C. brenneri* was found solely in sites impacted by humans, and never within the rainforest interior, suggesting that, like *C. elegans, C. brenneri* might experience dispersal benefits from human association. The lack of increasing species richness at larger spatial scales within French Guiana suggests that either i) we failed to adequately sample the full diversity of microhabitats accessible to *Caenorhabditis*, ii) the range of habitats and niche space for *Caenorhabditis* are relatively uniform across local and regional scales in this tropical region of South America, or iii) dispersal of *Caenorhabditis* is high enough in this region to render all if it accessible for colonization.

To explore the potential for substrate specificity in habitat use and colonization of new habitat patches, we conducted a collection of field surveys and experiments. *Caenorhabditis* were rarely encountered in leaf litter, but comprised the predominant group of nematodes in rotting fruit (2.7% of litter vs 20.6% of fruit samples had *Caenorhabditis). Caenorhabditis* also represented the dominant group that colonized fresh fruit baits, demonstrating that two of the most common species are capable of colonizing new resource patches rapidly (~2.5-7.5% baits/day; *C. nouraguensis* and *C. tropicalis*). Colonization of fresh resources occurred most readily when in close proximity to pre-existing native fruit patches on the rainforest floor. This analysis thus provides quantitative support for the received wisdom that *Caenorhabditis* occurs most frequently in high-nutrient rotting resources rich in active microbial growth, and that *Caenorhabditis* are not “soil nematodes” [10, 11, 14].

These observations raise the question of the most likely source of fresh resource patch founding: vector-mediated colonization or a “seed bank” of dauer larvae that crawl through the leaf litter. If primarily by vectors, as known for species like temperate-zone *C. japonica* [36], are flying versus crawling invertebrates primarily involved? Direct experimental tests are required to confidently assess these possibilities, though we are skeptical that *Caenorhabditis* colonists arrive primarily by crawling to new patches within the Nouragues Natural Reserve. A remaining further question is whether *Caenorhabditis* might colonize resource patches prior to fruits and flowers falling to the rainforest floor, for example, from pollinators or frugivorous animals.

Despite successful colonization of fresh resource patches by *Caenorhabditis*, many substrate patches were unoccupied and the rapid decay of rotting fruit substrates suggests that full exploitation of available resources by *Caenorhabditis* is likely to go unrealized due to imperfect colonization ability. Such dispersal limitation means that a single founder event per patch is likely, which raises several interesting issues. First, the high turnover of individual resource patches (<10 nematode generations duration), the hierarchical structuring of resource patches (individual fruits, individual source trees, landscape-scale clustering), and temporal dynamics in patch availability all make *Caenorhabditis* a tantalizing system for integrating metapopulation colonization-extinction processes with species co-existence [7, 10, 37, 38]. Second, priority effects for within-patch co-existence may be especially pronounced [39], also raising the potential for reproductive interference in accidental inter-species matings to interact with resource competition [40, 41]. Third, when individual vector transport (e.g. insects) constrains the number of colonist nematodes that colonize a new resource patch, such founder events could provide a recurring selective pressure favoring selfing as a means of reproductive assurance in the face of high metapopulation turnover [42].

*Caenorhabditis* life history strategies span the generalist to specialist spectrum, which is predicted to associate with population demography and genetic variation [43]. Distinct reproductive modes, self-fertilizing hermaphroditism and obligate outbreeding, also occur in *Caenorhabditis* that exert profound effects on genome evolution [43, 44]. In French Guiana, we find sympatric species spanning these traits and further find that the diverse *Caenorhabditis* community comprises a few abundant species and many numerically rare species. These attributes place these nematodes in a convenient position to test hypotheses about how organismal traits and population demography influence genetic variation and evolutionary change in the context of ecological community assembly [10, 43]. We anticipate that future studies of this and other hotspots of *Caenorhabditis* diversity will provide important further contributions to understanding the inter-relation of ecological factors, organismal features, and genomic trends in evolution.

## Declarations

**Ethics approval and consent to participate**: not applicable.

**Consent to publish**: not applicable.

**Availability of data and materials**:

*Partial ITS2 sequences of the two new species (*C. astrocarya* and *C. dolens*) will be deposited in GenBank (Additional File 9).

*This published work and the nomenclatural acts it contains have been registered in ZooBank, the online registration system for the ICZN. The ZooBank LSIDs (Life Science Identifiers) can be resolved and the associated information viewed through any standard web browser by appending the LSID to the prefix “http://zoobank.org/”. The LSID for this publication is urn:lsid:zoobank.org:pub:06DBDCA0-DD60-43BA-9A99-14F0748D4FE2. The ZooBank LSID for *Caenorhabditis dolens* is urn:lsid:zoobank.org:act:4D39DD5D-2120-42E5-8521-0F35D8710592 and the the LSID for *Caenorhabditis astrocarya* is urn:lsid:zoobank.org:act:EBE4FF82-F906-4237-9290-C84431F17D7D. The type culture specimens are deposited at the Caenorhabditis Genetics Center (<u>http://www.cbs.umn.edu/research/resources/cgc</u>).

*Live strain isolates are available from the authors.

*Collection data are provided as supplementary material in Additional Files 1-9).

## Authors’ contributions

CF and RS performed data collection and analysis in the lab. NC and RJ performed field experiments and collected data. AC and CB designed the experiments, collected and analyzed data, and wrote the manuscript. All authors read and approved the final manuscript.

## Acknowledgements

We thank the CNRS in French Guiana and its staff, in particular, Patrick cha□telet, Jéro□me Chave, Philippe Gaucher, Annaig Le Guen and Laetitia Proux. We further acknowledge support and advice provided by the Parc Amazonien de Guyane (Raphaëlle Rinaldo) and the Conseil régional de Guyane (Frédéric Blanchard). We thank Marie-Anne Félix for comments on the manuscript, Marine Guntz for help with sampling and Anne Vielle, Clotilde Gimond, Paul Vigne and Emilie Demoinet for help with freezing numerous *Caenorhabditis* isolates.

